# DLDN-Bench: A Benchmark Framework for Deep Learning De Novo Peptide Sequencing in Proteomics

**DOI:** 10.64898/2026.06.10.728383

**Authors:** Jannik Schneider, Sonja Hartwig, Alexandra Chadt, Stefan Lehr, Hadi Al-Hasani, Michael Turewicz

## Abstract

De novo peptide sequencing is an essential approach for analyzing mass spectrometry data because it enables the identification of novel peptides without relying on protein sequence databases. Recent advances in deep learning have substantially improved the performance of de novo sequencing methods, but the rapid emergence of new models has led to heterogeneous evaluation practices and limited comparability. To address this, we introduce DLDN-Bench, a benchmark framework including a set of benchmark datasets derived from human muscle biopsy mass spectrometry data retrieved from PRIDE and annotated through consensus across multiple widely used database search engines. Using these datasets, we systematically benchmark recent deep learning-based de novo sequencing tools alongside traditional approaches. Performance is assessed using established metrics, including precision and coverage relative to a pseudo-ground truth defined by cross-engine agreement. To demonstrate the utility of DLDN-Bench, we benchmark four recent deep learning models and make all results publicly available. This benchmark framework provides a standardized basis for comparing state-of-the-art methods and offers an extensible resource for evaluating future tools in de novo peptide sequencing.

**Code availability:** https://github.com/ddz-icb/DLDN-Bench

**Data availability:** https://doi.org/10.5281/zenodo.19627459

## Introduction

Bottom-up proteomics is a widely used strategy in which proteins are enzymatically digested into peptides that are separated by liquid chromatography and analyzed by tandem mass spectrometry (LC-MS/MS). The resulting MS/MS spectra contain mass-to-charge values and fragmentation patterns that enable peptide identification. Three major approaches are commonly used: database searching, spectral library searching, and de novo sequencing.

In database searches, experimental spectra are matched against theoretical spectra generated from protein sequence databases such as UniProt (UniProt Consortium 2025). Search engines including MSGF+ (Kim and Pevzner 2014), Comet (Eng, Jahan, and Hoopmann 2013), and MSFragger (Kong *et al*. 2017) compare observed and predicted fragmentation patterns to identify peptides and localize post-translational modifications. Spectral library searches instead compare experimental spectra to previously annotated spectra in curated libraries, often improving confidence when high-quality reference spectra are available.

De novo peptide sequencing infers amino acid sequences directly from MS/MS spectra without relying on a protein database. This makes it particularly valuable for identifying novel peptides, unexpected modifications, or sequences from organisms with incomplete or unavailable reference proteomes. Since the introduction of DeepNovo (Tran *et al*. 2017) in 2017, the field has been transformed by deep learning-based models, which leverage large annotated datasets to learn the mapping from spectra to peptide sequences and have achieved substantial performance improvements (Bittremieux *et al*. 2026).

To address the need for standardized and reproducible evaluation of modern deep learning– based de novo sequencing methods, we introduce DLDN-Bench, a benchmark framework that provides curated datasets, unified evaluation metrics, and an extensible workflow for comparing state-of-the-art and future tools, including a demonstration benchmarking of four recent deep learning models whose results are made publicly available.

## Methods

### Datasets

Raw LC–MS/MS data were obtained from the PRIDE repository (Deshmukh and Mann 2020, Kenny *et al*. 2020, Dreher *et al*. 2024, Turewicz *et al*. 2025) (accessions: PXD043425, PXD006882, PXD043200, PXD012824) and consisted of human skeletal muscle biopsy samples collected under different physiological conditions, including insulin-treated and untreated states. The protein composition of muscle tissue poses a challenge for identification algorithms because muscle contains a large number of known high-molecular-weight proteins, many of which share substantial sequence similarity. As a result, proteomic datasets derived from muscle samples are particularly suitable for benchmarking identification methods. All raw files were converted to MGF format using msconvert (ProteoWizard) (Adusumilli and Mallick 2017) with centroiding enabled via the peak picking filter. The overall dataset selection and preprocessing workflow is summarized in Fig. 1a and Supplemental Table 2.

**Figure 1.**
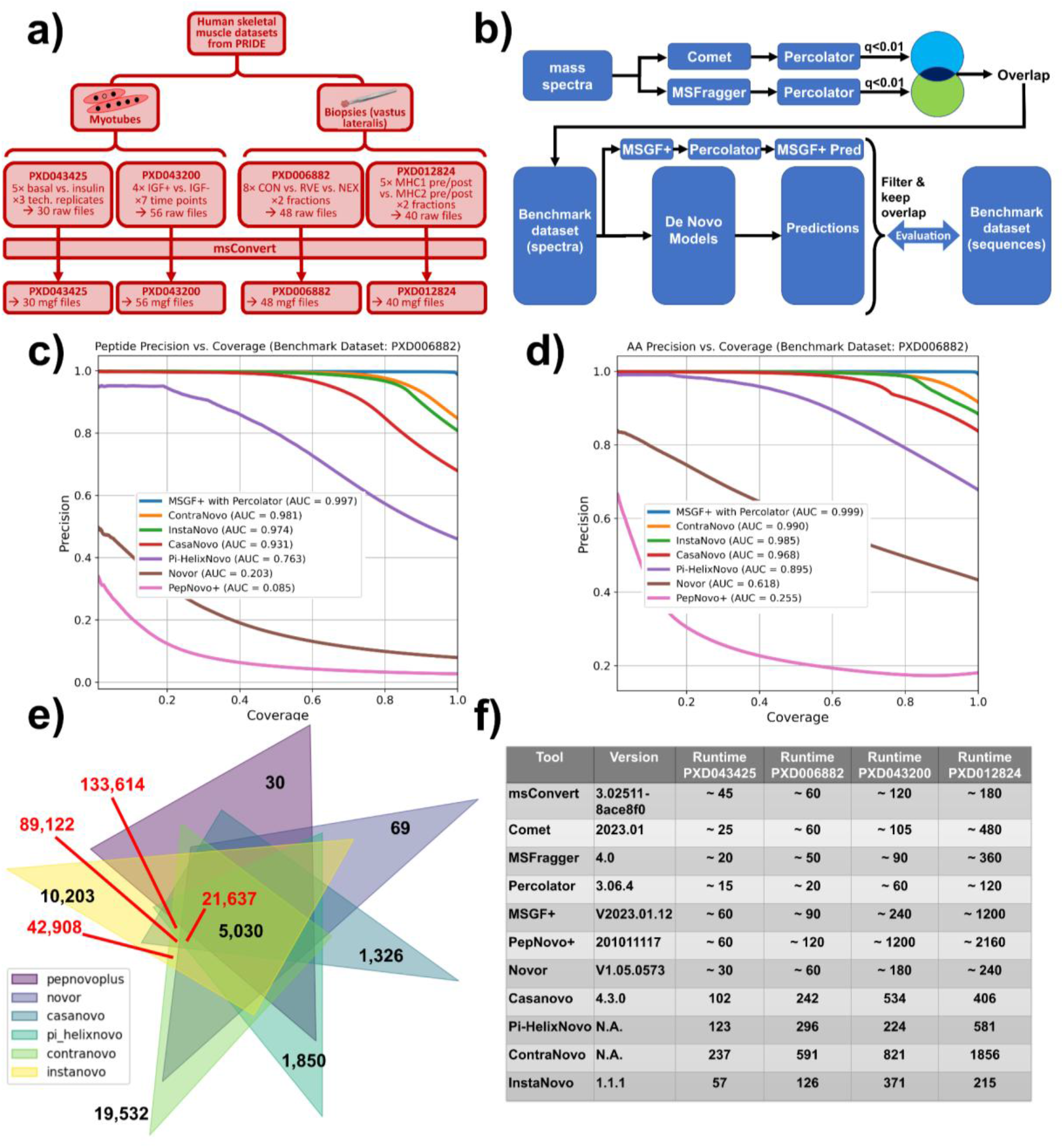
Benchmark dataset, workflow and results. **a)** Benchmark datasets. **b)** Comparison workflow. **c)** Peptide precision vs. coverage results for one benchmark dataset and multiple traditional and deep learning tools. **d)** Amino acid precision vs. coverage results for the same dataset and tools. **e)** Venn diagram of true positive predictions. The diagram shows the overlap of correctly identified peptide sequences among the evaluated tools, based on matches to the pseudo-ground truth. ContraNovo and InstaNovo exhibit the largest sets of unique true positives, indicating complementary strengths relative to the other methods. **f)** Runtimes of benchmarked tools in minutes.

### Benchmark dataset generation

Each dataset was searched using two independent database search engines to match the centroided MGF spectra against theoretical peptide spectra. Searches were performed with a standard set of modifications: carbamidomethylation of cysteine as a fixed modification, and oxidation of methionine, N-terminal methionine loss, and N-terminal acetylation as variable modifications.

To obtain high-confidence peptide–spectrum matches (PSMs), a two-step filtering strategy was applied:

1. Rescoring and FDR control: Search results from each engine were rescored using Percolator (The *et al*. 2016) and filtered at a q-value < 0.01, following standard proteomics practice for controlling the false discovery rate.
2. Consensus filtering: Only PSMs identified by both search engines after FDR filtering were retained. This intersection yields a high-confidence set of consensus PSMs that serves as the pseudo-ground truth for benchmarking.

### Traditional and deep learning-based tools

The de novo peptide sequencing tools were grouped into traditional and deep learning-based approaches. Traditional tools included PepNovo+ (Frank and Pevzner 2005) and Novor (Ma 2015), both executed through the command-line interface of DeNovoGUI (Muth *et al*. 2014).

For the deep learning-based category, we selected recent state-of-the-art models for which pretrained weights were publicly available and trained on data dependent acquisition (DDA) spectra. The evaluated models were CasaNovo (Yilmaz *et al*. 2022), Pi-HelixNovo (Yang *et* al. 2024), ContraNovo (Jin *et al*. 2023), and InstaNovo (Eloff *et al*. 2025).

In addition to de novo tools, MS-GF+ was applied to the benchmark datasets, followed by rescoring with Percolator, to provide a database-search-based reference point for comparison.

### Evaluation Criteria

The predicted peptide sequences were evaluated using precision and coverage at both the peptide level and the amino-acid level, following the definitions commonly used in recent deep learning-based de novo sequencing studies (Yilmaz *et al*. 2022). Precision measures the proportion of correctly predicted peptides (or amino acids) among all predictions made at a given confidence threshold, while coverage quantifies the proportion of the pseudo-ground-truth peptides (or amino acids) that are recovered by the model.

To visualize the trade-off between these metrics, precision-coverage curves were generated for each benchmark dataset and for both evaluation levels. The area under the curve (AUC) was computed to summarize overall performance across confidence thresholds, enabling direct comparison between tools.

### Resources

All steps related to benchmark dataset generation and the execution of the traditional de novo sequencing tools were performed on the same local workstation (Intel Core Ultra 7 155H with 1.4GHz, 64 GB RAM) to ensure consistent runtime conditions. These were executed in the Windows Subsystem for Linux (WSL) environment. In contrast, the deep learning–based models were executed in a GPU-accelerated cloud environment (Google Colab; 50 GB RAM; NVIDIA A100, 48 GB VRAM), as these tools require substantial GPU resources for efficient inference.

## Results

As a central outcome of this work, we provide DLDN-Bench, an open and extensible benchmark framework available to the community through GitHub. DLDN-Bench enables reproducible evaluation of both the recent deep learning–based de novo sequencing models included in this study and additional models contributed by users, using the benchmark datasets described in this work. The framework also supports the integration of new annotated MS/MS datasets, allowing users to expand the benchmark beyond the provided data. To demonstrate the utility of DLDN-Bench as an unbiased, independent, and transparent evaluation resource, we performed a comprehensive benchmarking of four recent deep learning models and make these results publicly available.

Across the evaluated PRIDE dataset, ContraNovo and InstaNovo achieved the highest de novo sequencing performance among the tested tools. In the peptide-level precision– coverage analysis (Fig. 1b), ContraNovo reached an AUC of 0.981 and InstaNovo an AUC of 0.974, approaching the performance of the database-search reference MS-GF+ (AUC = 0.997). The traditional tools performed substantially worse, with Novor achieving an AUC of 0.203 and PepNovo+ only 0.085.

A similar pattern was observed at the amino-acid level (Fig. 1d). ContraNovo (AUC = 0.990) and InstaNovo (AUC = 0.985) again nearly matched MS-GF+ (AUC = 0.999), while the traditional tools showed markedly lower performance (Novor AUC = 0.618; PepNovo+ AUC = 0.255).

The overlap analysis of correctly identified peptide sequences (Fig. 1e) revealed that ContraNovo and InstaNovo each contributed a large number of unique true positives (ContraNovo: 19,532; InstaNovo: 10,203). Their pairwise overlap was also substantial, with 42,908 peptides identified exclusively by both deep learning models, indicating complementary strengths relative to the other tools.

Runtime comparisons (Fig. 1f) showed the opposite trend: the traditional tools were the fastest, while the deep learning–based methods required longer inference times. Among the deep learning models, InstaNovo exhibited shorter runtimes than ContraNovo.

Results for the remaining benchmark datasets followed the same overall pattern (see Supplemental Figures 1-12). Overall, our evaluation confirms the substantial performance gains reported in previous studies of recent deep learning–based de novo sequencing tools (Yilmaz et al. 2022, Jin *et al*. 2023, Yang et al. 2024, Eloff *et al*. 2025, Van Den Bossche *et al*. 2025).

## Discussion

Applying MS-GF+ to the benchmark datasets provides a useful reference point that approximates a gold-standard scenario in which an appropriate protein database is available. This comparison also relates to the ongoing question of whether modern de novo sequencing tools can serve as standalone alternatives for peptide identification (Bittremieux *et al*. 2026). Our results show that the best-performing deep learning models, ContraNovo and InstaNovo, approach MS-GF+ performance up to a coverage of approximately 70%. This suggests that, when lower-confidence predictions are excluded, these tools can indeed function as standalone identification methods in settings where database searches are not feasible. Nevertheless, for maximizing peptide recovery, hybrid strategies that combine database searching with subsequent de novo analysis are likely to remain advantageous.

The evaluation pipeline presented here is designed to be extensible. New de novo sequencing tools can be incorporated easily and benchmarked against the provided datasets, and users may also supply additional annotated datasets to broaden the evaluation space. By offering standardized datasets, metrics, and workflows, this benchmark aims to support the community’s call for more independent and reproducible evaluations in light of the rapidly growing number of deep learning–based de novo peptide sequencing methods.

