## Supplemental for "DLDN-Bench: A Benchmark Framework for Deep Learning De Novo Peptide Sequencing in Proteomics"

### Supplemental Figures

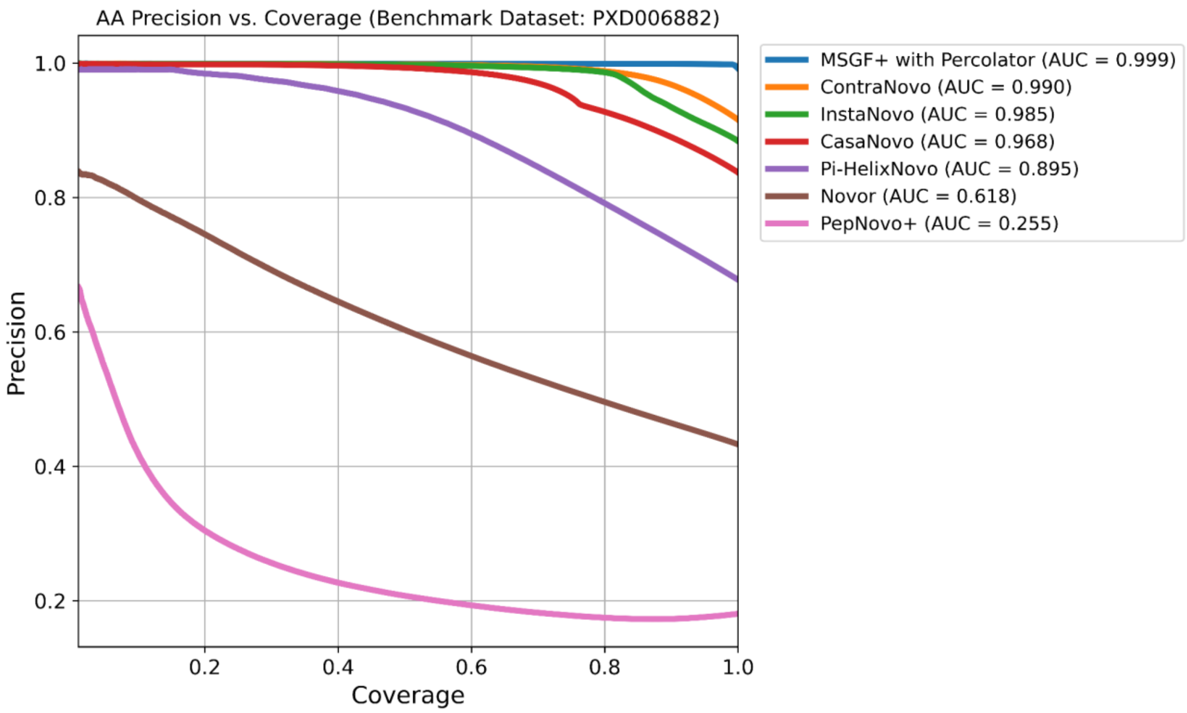


***Supplemental Figure 1: Amino acid precision vs. coverage results for PXD006882.*** *The Precision-Coverage curves on the amino acid level on the benchmark dataset PXD006882. MSGF+ with Percolator is shown for comparison as gold standard since the ground truth database was provided. The area-under-the-curve (AUC) can be interpreted as the probability that the model assigns a higher score/confidence to a true-positive amino acid. ContraNovo(AUC=0.990) and InstaNovo(AUC=0.985) almost achieve similar results than the database search with MSGF+ without having the database provided.*

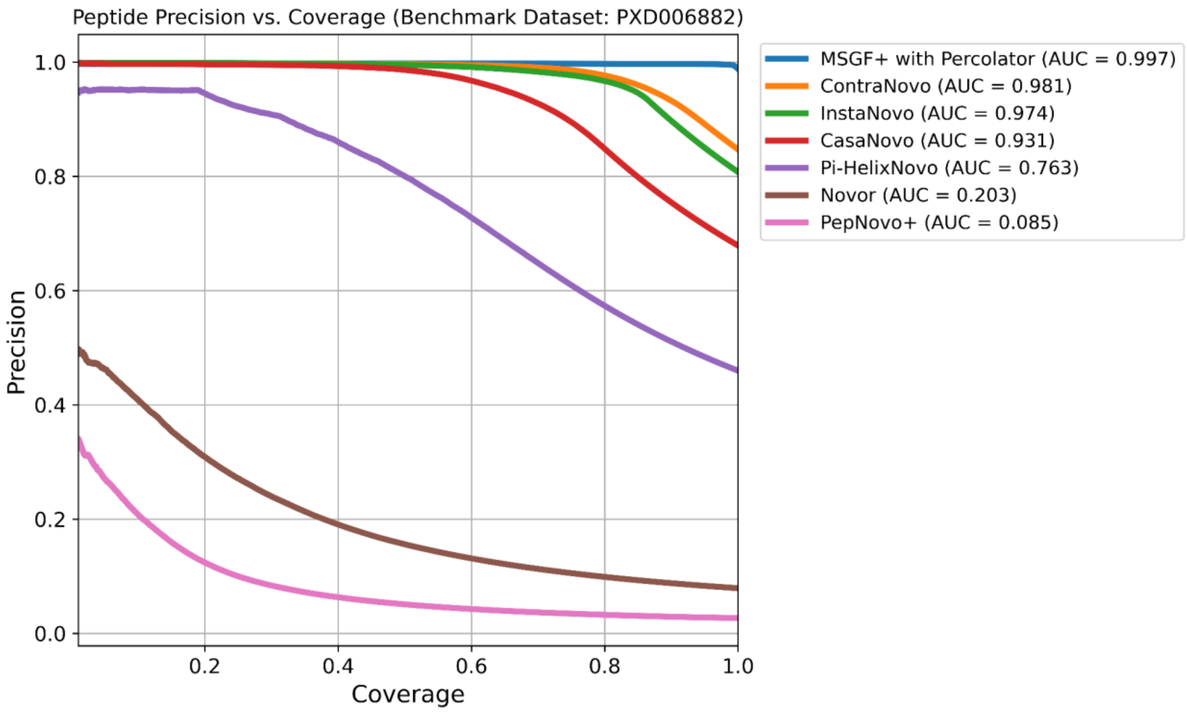


***Supplemental Figure 2: Peptide precision vs. coverage results for PXD006882.*** *The Precision-Coverage curves on the peptide level on the benchmark dataset PXD006882. The ranking is similar as in supplemental figure 1, however slightly below the amino acid precision since predicting the whole peptide sequence correct is the harder task.*

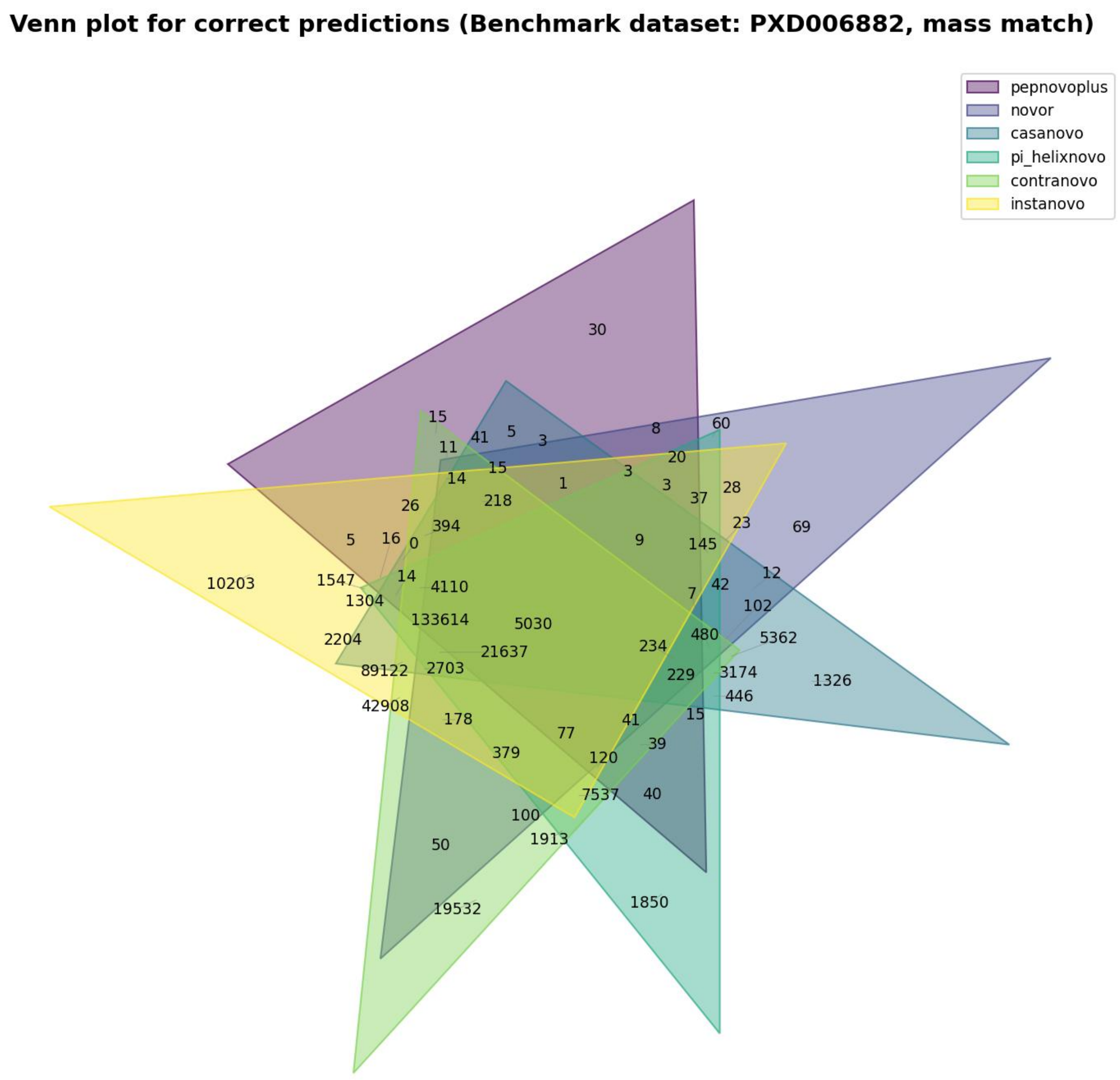


***Supplemental Figure 3: Venn diagram of correct predictions for PXD006882.*** *The Venn plot shows the overlaps of the peptide sequence predictions between the models on the benchmark dataset PXD006882. ContraNovo achieves the largest unique set (19532) which aligns with the precision-coverage performance. The largest set in general is the overlap of all models (133614).*

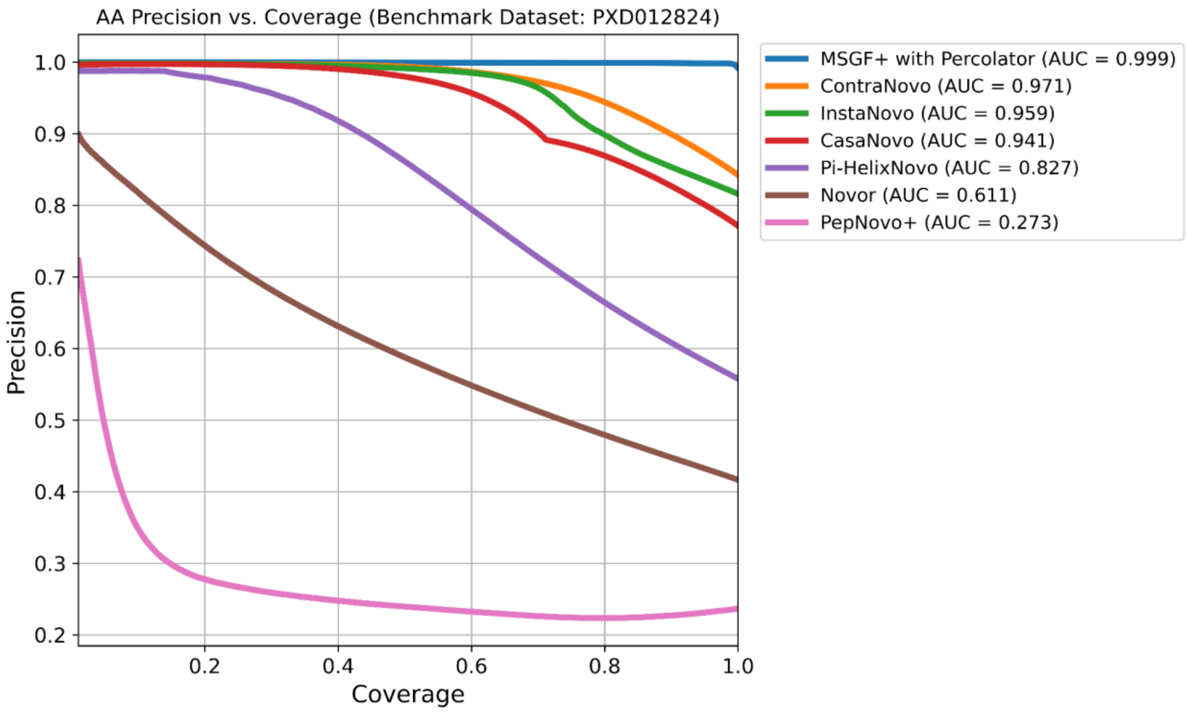


***Supplemental Figure 4: Amino acid precision vs. coverage results for PXD012824.*** *The Precision-Coverage curves on the amino acid level on the benchmark dataset PXD012824. The ranking remains similar as in supplementary figures 1-2, however the gap between ContraNovo(AUC=0.971) and InstaNovo(AUC=0.959) is slightly larger.*

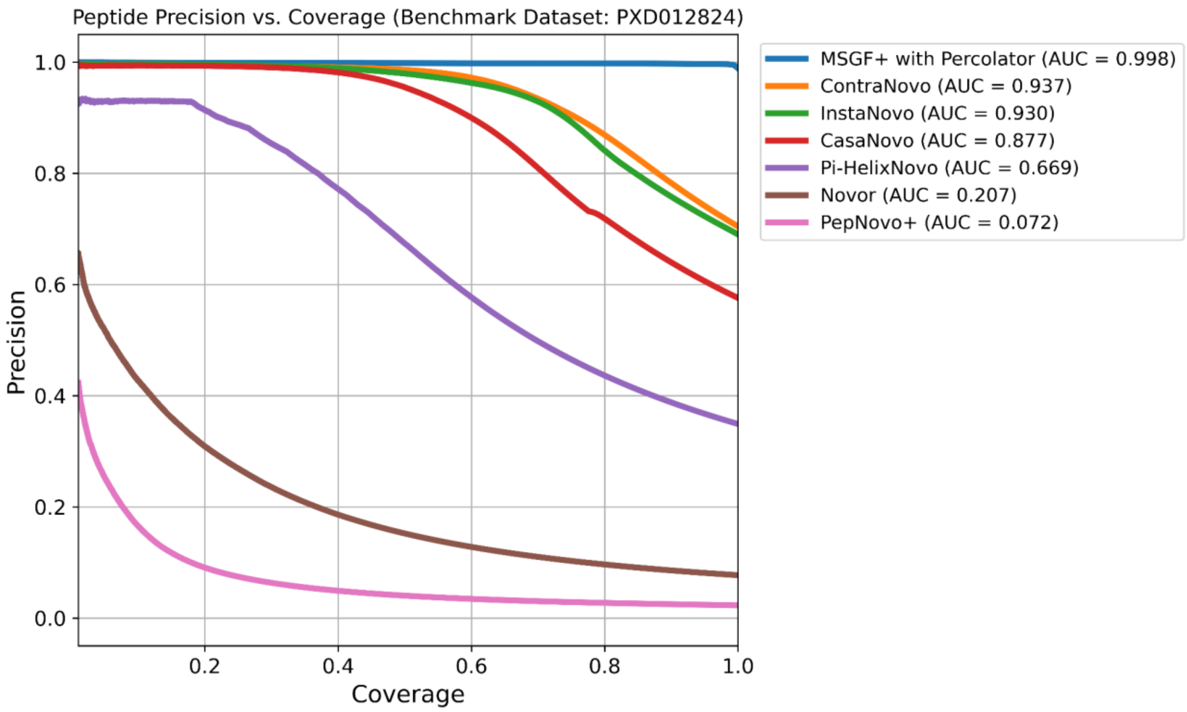


***Supplemental Figure 5: Peptide precision vs. coverage results for PXD012824.*** *The Precision-Coverage curves on the peptide level on the benchmark dataset PXD012824. The ranking remains similar as in supplementary figures 1-2.*

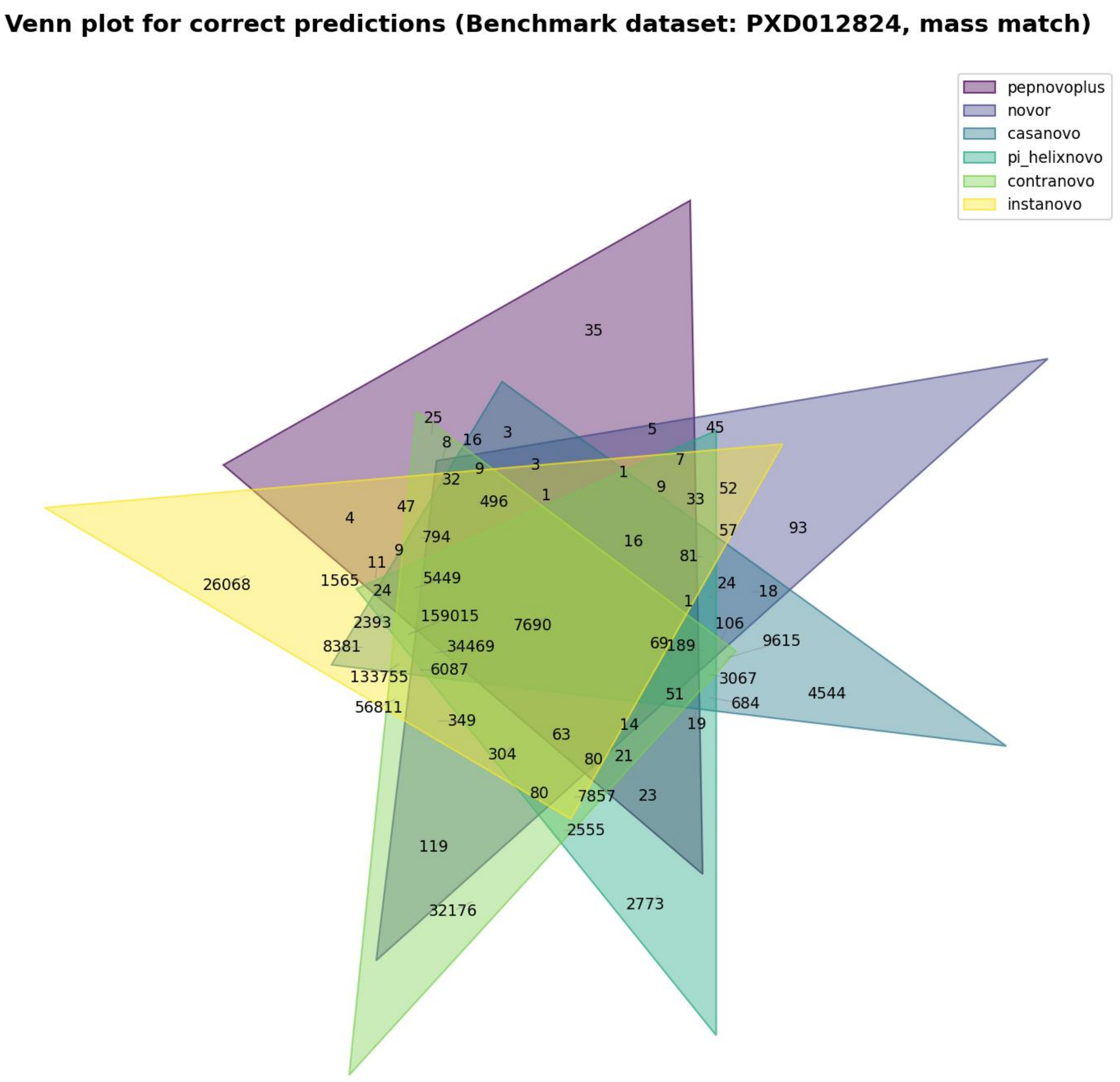


***Supplemental Figure 6: Venn diagram of correct predictions for PXD012824.*** *The Venn plot shows the overlaps of the peptide sequence predictions between the models on the benchmark dataset PXD012824. ContraNovo achieves the largest unique set (32176) which aligns with the precision-coverage performance. The largest set in general is the overlap of all models (159015).*

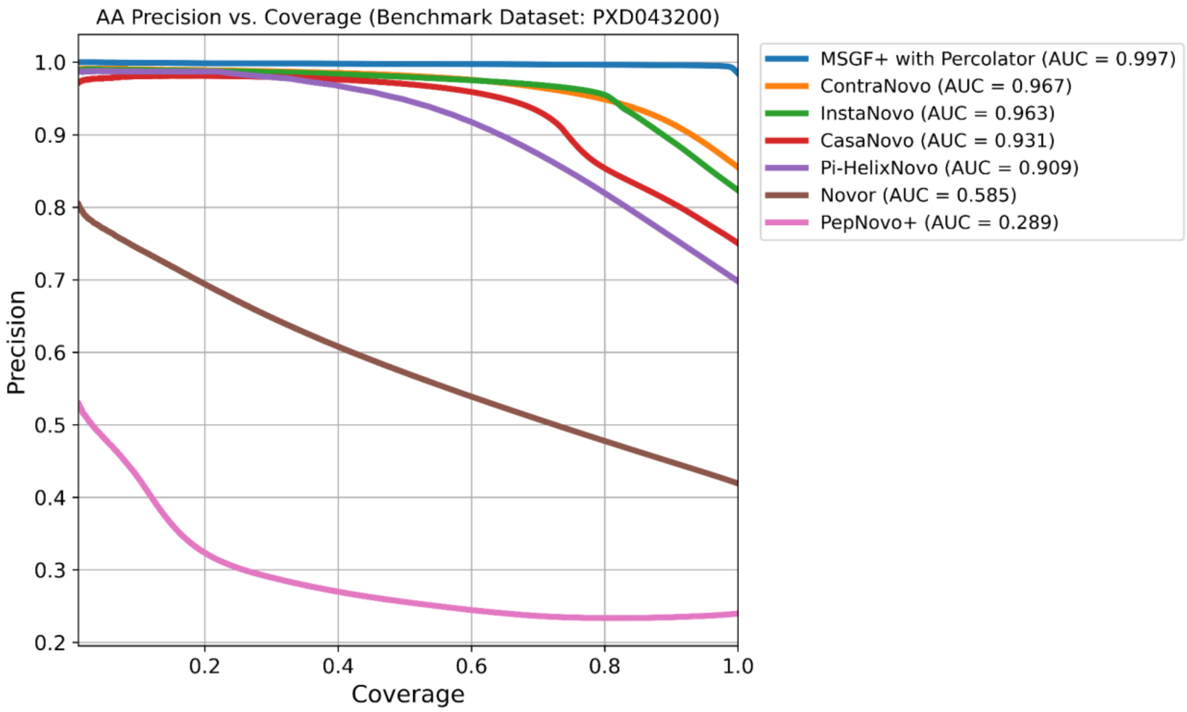


***Supplemental Figure 7: Amino acid precision vs. coverage results for PXD043200.*** *The Precision-Coverage curves on the amino acid level on the benchmark dataset PXD043200. The ranking remains similar as in supplementary figures 1-2.*

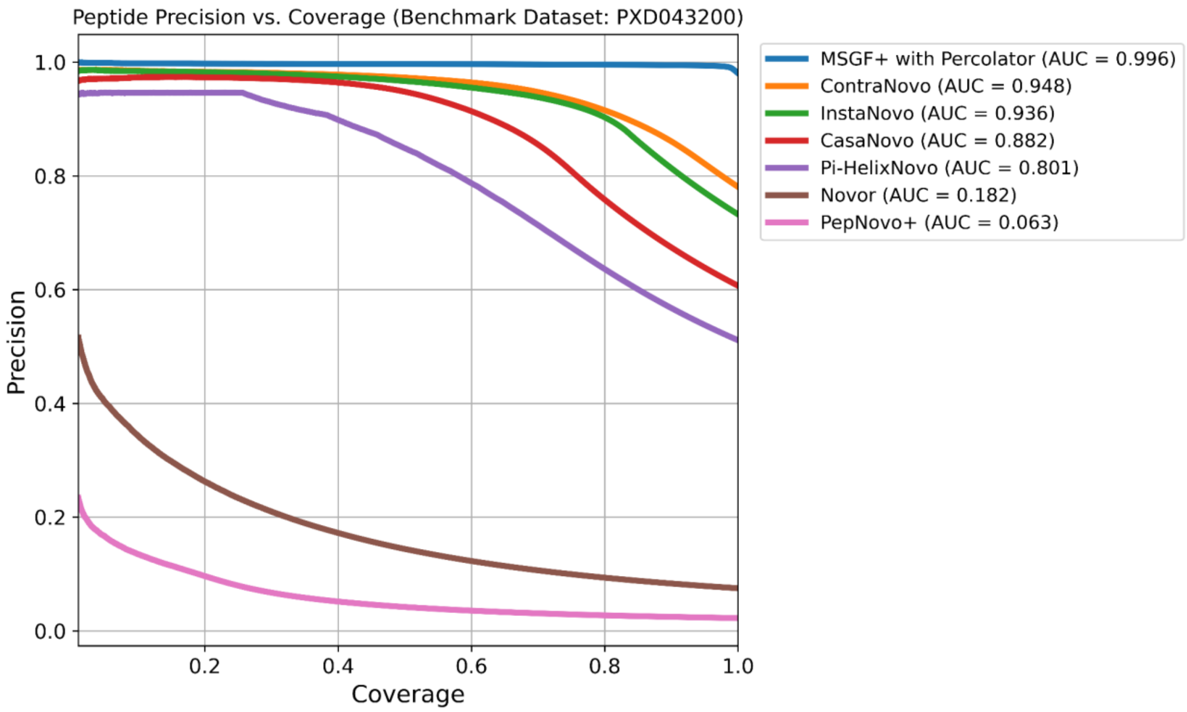


***Supplemental Figure 8: Peptide precision vs. coverage results for PXD043200.*** *The Precision-Coverage curves on the peptide level on the benchmark dataset PXD043200. The ranking remains similar as in supplementary figures 1-2.*

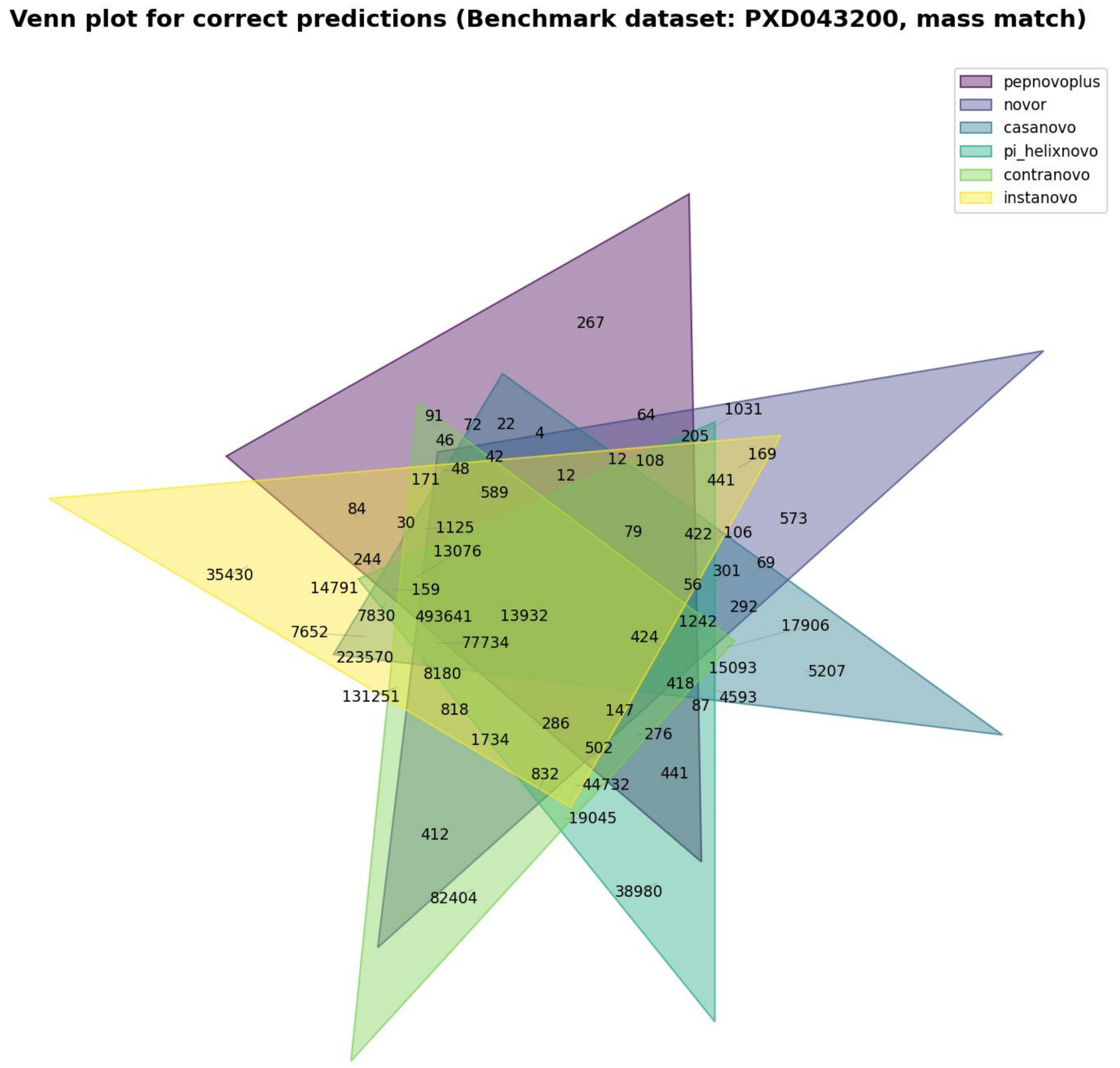


***Supplemental Figure 9: Venn diagram of correct predictions for PXD043200.*** *The Venn plot shows the overlaps of the peptide sequence predictions between the models on the benchmark dataset PXD043200. ContraNovo achieves the largest unique set (82404) which aligns with the precision-coverage performance. The largest set in general is the overlap of all models (493641).*

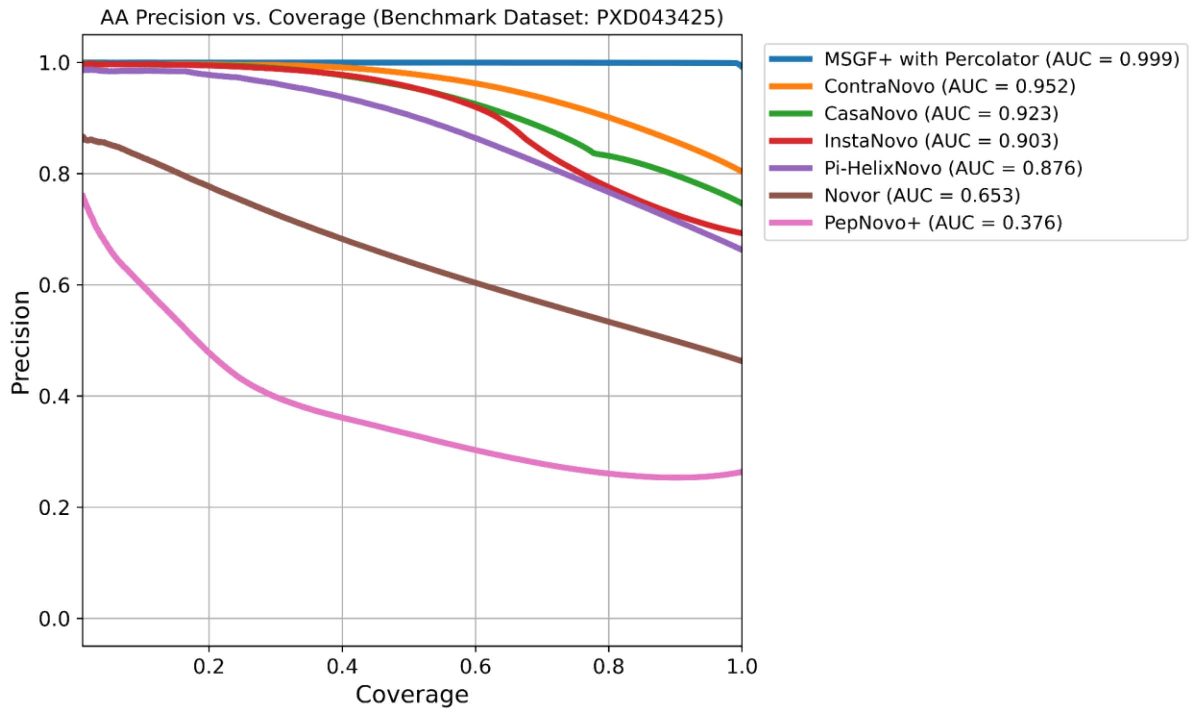


***Supplemental Figure 10: Amino acid precision vs. coverage results for PXD043425.*** *The Precision-Coverage curves on the amino acid level on the benchmark dataset PXD043425. The ranking remains similar as in supplementary figures 1-2.*

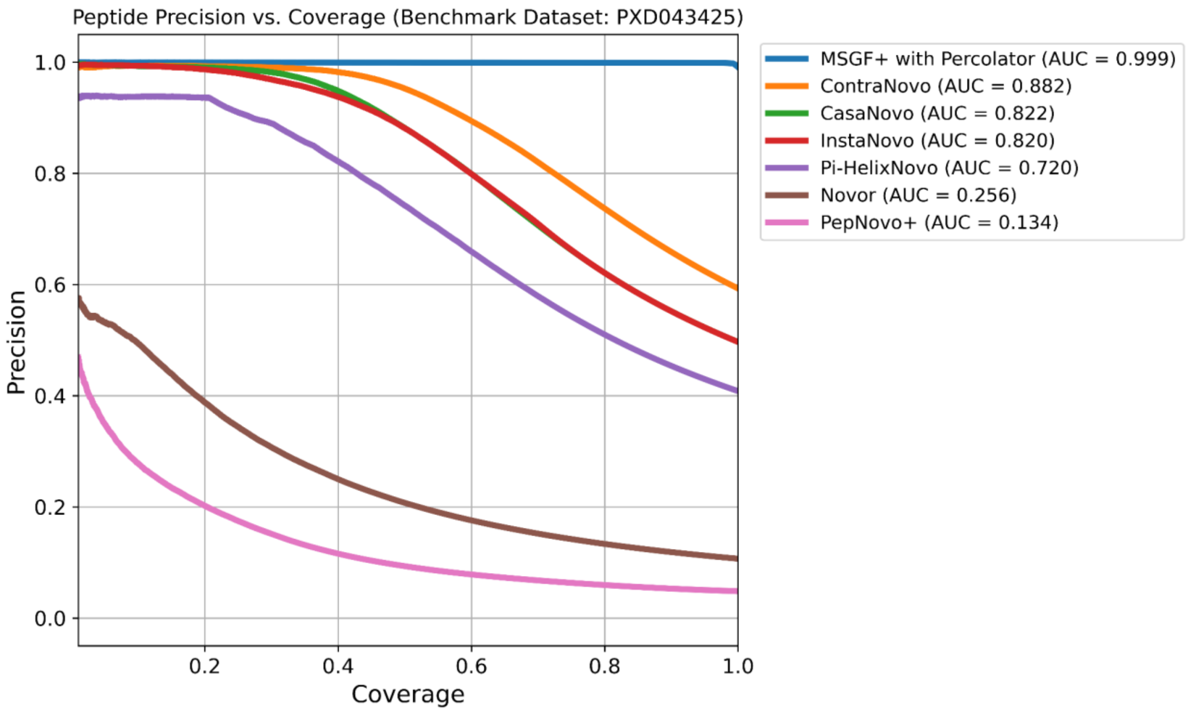


***Supplemental Figure 11: Peptide precision vs. coverage results for PXD043425.*** *The Precision-Coverage curves on the peptide level on the benchmark dataset PXD043425. The ranking remains similar as in supplementary figures 1-2 but generally below the performance on the other benchmark datasets.*

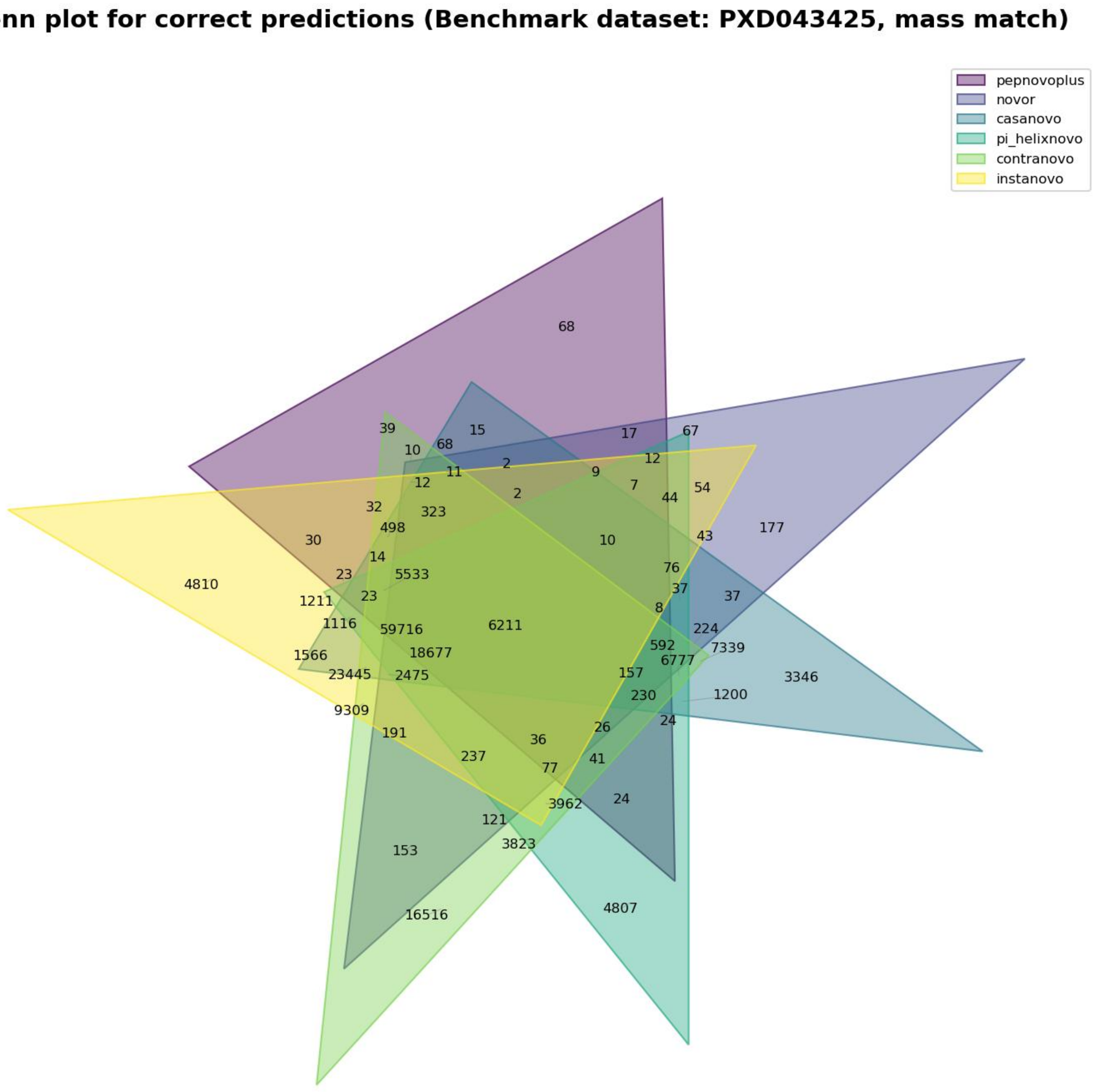


***Supplemental Figure 12: Venn diagram of correct predictions for PXD043425.*** *The Venn plot shows the overlaps of the peptide sequence predictions between the models on the benchmark dataset PXD043425. ContraNovo achieves the largest unique set (16516) which aligns with the precision-coverage performance. The largest set in general is the overlap of all models (59716).*

### Supplemental Tables

| Component | Google Colab (Pro) | Local Machine (WSL) |
|---|---|---|
| CPU | Intel Xeon CPU @ 2.20GHz (virtualized) | Intel® Core™ Ultra 7 155H, 1400MHz, 16 Core(s), 22 logical core(s) |
| RAM | 50GB | 64GB |
| GPU | L4 GPU | Not used |
| Operating System | Ubuntu 20.04 LTS | Ubuntu 22.04 (WSL) |

***Supplemental Table 1: Hardware specifications of the utilized systems.*** *The specifications of the environment (Google Colab) which was used to execute the deep learning based tools as well as the local machine (WSL) which was used for the execution of the preprocessing and database search engines.*

| Benchmark dataset (Pride accession) | Link | # Peptide-spectrum matches (after workflow) |
|---|---|---:|
| PXD006882 | https://www.ebi.ac.uk/pride/archive/projects/PXD006882 | 400.625 |
| PXD012824 | https://www.ebi.ac.uk/pride/archive/projects/PXD012824 | 654.984 |
| PXD043200 | https://www.ebi.ac.uk/pride/archive/projects/PXD043200 | 1.473.250 |
| PXD043425 | https://www.ebi.ac.uk/pride/archive/projects/PXD043425 | 281.404 |

***Supplemental Table 2: Benchmark datasets statistics.*** *The four datasets which were fetched from the PRIDE archive along with the number of remaining peptide-spectrum matches after the described workflow which were used for the evaluation.*
